# Analysis of Rapidly Emerging Variants in Structured Regions of the SARS-CoV-2 Genome

**DOI:** 10.1101/2020.05.27.120105

**Authors:** Sean P. Ryder, Brittany R. Morgan, Francesca Massi

## Abstract

The severe acute respiratory syndrome coronavirus 2 (SARS-CoV-2) pandemic has motivated a widespread effort to understand its epidemiology and pathogenic mechanisms. Modern high-throughput sequencing technology has led to the deposition of vast numbers of SARS-CoV-2 genome sequences in curated repositories, which have been useful in mapping the spread of the virus around the globe. They also provide a unique opportunity to observe virus evolution in real time. Here, we evaluate two cohorts of SARS-CoV-2 genomic sequences to identify rapidly emerging variants within structured cis-regulatory elements of the SARS-CoV-2 genome. Overall, twenty variants are present at a minor allele frequency of at least 0.5%. Several enhance the stability of Stem Loop 1 in the 5’UTR, including a set of co-occurring variants that extend its length. One appears to modulate the stability of the frameshifting pseudoknot between ORF1a and ORF1b, and another perturbs a bi-stable molecular switch in the 3’UTR. Finally, five variants destabilize structured elements within the 3’UTR hypervariable region, including the S2M stem loop, raising questions as to the functional relevance of these structures in viral replication. Two of the most abundant variants appear to be caused by RNA editing, suggesting host-viral defense contributes to SARS-CoV-2 genome heterogeneity. This analysis has implications for the development of therapeutics that target viral cis-regulatory RNA structures or sequences, as rapidly emerging variations in these regions could lead to drug resistance.

## Introduction

The betacoronaviridae are non-segmented single-stranded positive sense viruses with an RNA genome of approximately thirty kilobases in length. This family poses a significant threat to human health. In addition to causing approximately 30% of annual upper respiratory infections (Stadler et al. 2003; Su et al. 2016), it is responsible for three major outbreaks of severe acute respiratory syndrome (SARS, MERS, and COVID-19) since the turn of the century (Drosten et al. 2003; Ksiazek et al. 2003; Zhong et al. 2003; Zaki et al. 2012; Wang et al. 2020). COVID-19 is a unique form of pneumonia characterized by high fever, dry cough, and occasionally catastrophic hypoxia. It was first described in the city of Wuhan, Huibei Province, in the fall of 2019 (Chan et al. 2020; Li et al. 2020; Wang et al. 2020). A novel virus termed severe acute respiratory syndrome coronavirus 2 (SARS-CoV-2) was identified as the cause of this disease (Wang et al. 2020; Wu et al. 2020). The rapid spread of the virus led to a global pandemic that caused significant morbidity and mortality and disruption of daily life for millions of people. The extraordinary impact of this virus fueled strong interest in understanding its pathophysiology and epidemiology with the hope of developing new treatments and approaches to limit its spread.

The SARS-CoV-2 infection cycle is similar to that of other betacoronaviridae (Zheng 2020). Following attachment of the virus to the host cell and membrane fusion, viral genomic RNA is introduced to the host cell where it is translated to produce a polyprotein encoding the viral replicase, proteases, and several accessory proteins. The replicase is an RNA-dependent RNA polymerase that produces full-length antigenomic sequence that serves as a template for the production of additional copies of the viral genome and several nested subgenomic RNAs (sgRNAs) that encode the structural components of the virion.

Conserved stem loop structures are present in both coding and noncoding regions the SARS-CoV-2 RNA genome (Ramya Rangan 2020). They cluster in the 5’UTR, the N-terminal portion of ORF-1a, at the junction of ORF1a and ORF1b, and in the 3’UTR. While their precise role is not known, their function can be inferred from studies of related elements in mouse hepatitis virus (MHV) and other coronaviridae (Yang and Leibowitz 2015; Madhugiri et al. 2016). The structured elements have regulatory roles in various aspects of viral replication, sgRNA synthesis, and translation. Though they are divergent in sequence, the structures appear to be conserved, and in some cases elements from SARS-CoV can functionally substitute for those in MHV with little impact on viral replication. (Goebel et al. 2004; Kang et al. 2006a; Kang et al. 2006b; Zust et al. 2008; Chen and Olsthoorn 2010; Yang and Leibowitz 2015).

DNA sequencing technology has progressed remarkably since the SARS outbreak of 2003 (Geoghegan and Holmes 2018; Zhang et al. 2018). It is now routine to determine the sequence of the ~30 kilobase viral genome using high throughput sequencing technology (Wu et al. 2020). As a result, scientists and medical professionals from around world have sequenced the SARS-CoV-2 genome from patient isolates and disseminated their findings through data repositories (e.g. the GISAID EpiCoV database) at unprecedented speed (Elbe and Buckland-Merrett 2017; Shu and McCauley 2017; Coronaviridae Study Group of the International Committee on Taxonomy of 2020; Wu et al. 2020). This has enabled the construction of molecular phylogenies that have guided our understanding of the virus transmission history, its basal mutation rate, and its potential to evade emerging therapeutics and vaccines (Bedford et al. 2020; Chu et al. 2020; Forster et al. 2020; Kim et al. 2020b; Lv et al. 2020; Pachetti et al. 2020; Pinto et al. 2020). At the time of this writing, almost 25,000 SARS-CoV-2 genome sequences have been deposited in the GISAID EpiCoV database (www.gisaid.org) and are available through a database access agreement (Elbe and Buckland-Merrett 2017; Shu and McCauley 2017). Over 3500 SARS-CoV-2 genome sequences have been deposited into the National Center for Biotechnology Information (NCBI) Genbank (ncbi.nih.nlv.gov/genbank/sars-cov-2-seqs) and are freely available to the public.

Here, we analyzed both cohorts to identify and characterize rapidly emerging variations within the cis-regulatory RNA structures of the virus genome. Our analysis reveals twenty rapidly emerging variants including two that likely arose through RNA editing. The data identify SL1 of the 5’UTR as a hot spot for viral mutation, where most mutations stabilize the stem loop structure. The data also show that structured elements in the 3’UTR hypervariable region, including the enigmatic S2M loop, contain rapidly emerging variations predicted to be destabilizing. The results provide insight into the relevance of the proposed viral RNA structures, and present a roadmap to avoid potential confounds to RNA therapeutic development.

## Results

### Identification of rapidly emerging variants in structured regions of the SARS-CoV-2 genome

The genome sequence for viral isolate Wuhan-Hu-1 (Genbank MN908947) was used as a reference genome (Wu et al. 2020). The 5’UTR (1–265), the structured region of ORF1a (266–450), the frameshifting pseudoknot (13,457–13,546), or the 3’UTR (29,543–29,903) were used as queries in a BLASTn search of the NCBI Betacoronavirus database filtered for SARS-CoV-2 (Altschul et al. 1990; Camacho et al. 2009). An average of 3600 ± 160 hits were recovered from each query. The sequences recovered from BLASTn were aligned with MAFFT using the FFT-NS-2 algorithm to produce a multiple sequence alignment (MSA) (Katoh et al. 2002). The MSA was then input into WebLogo 3 to calculate the positional occupancy, entropy, and allele frequency for each query (Supplementary Table 1) (Crooks et al. 2004). The occupancy defines the number of A, C, G, or U bases observed at each position (denoted as weight in the WebLogo3 output), the entropy defines the positional information content (lower value equals more variation), and the allele frequency defines the fractional occupancy of each nucleotide at each position. The results reveal high occupancy (>90%) from position 57 of the 5’UTR through position 29,836 of the 3’UTR, but the occupancy drops off significantly near the 5’ and 3’ ends of the genome (Fig. 1A), dipping below 20%. This is presumably due to difficulty of capturing the ends of the genome in sequencing library production. Nevertheless, even the extreme termini have coverage of more than 300 genomes. The positional entropy scores identify multiple variations in both low and high occupancy regions suggesting that variant entropy is not overly skewed by the terminal deficiencies in the genomic sequencing data. In total, fourteen variants with a minor allele frequency (MAF) of greater than 0.005 (0.5%) were identified by this approach.

**Figure 1.**
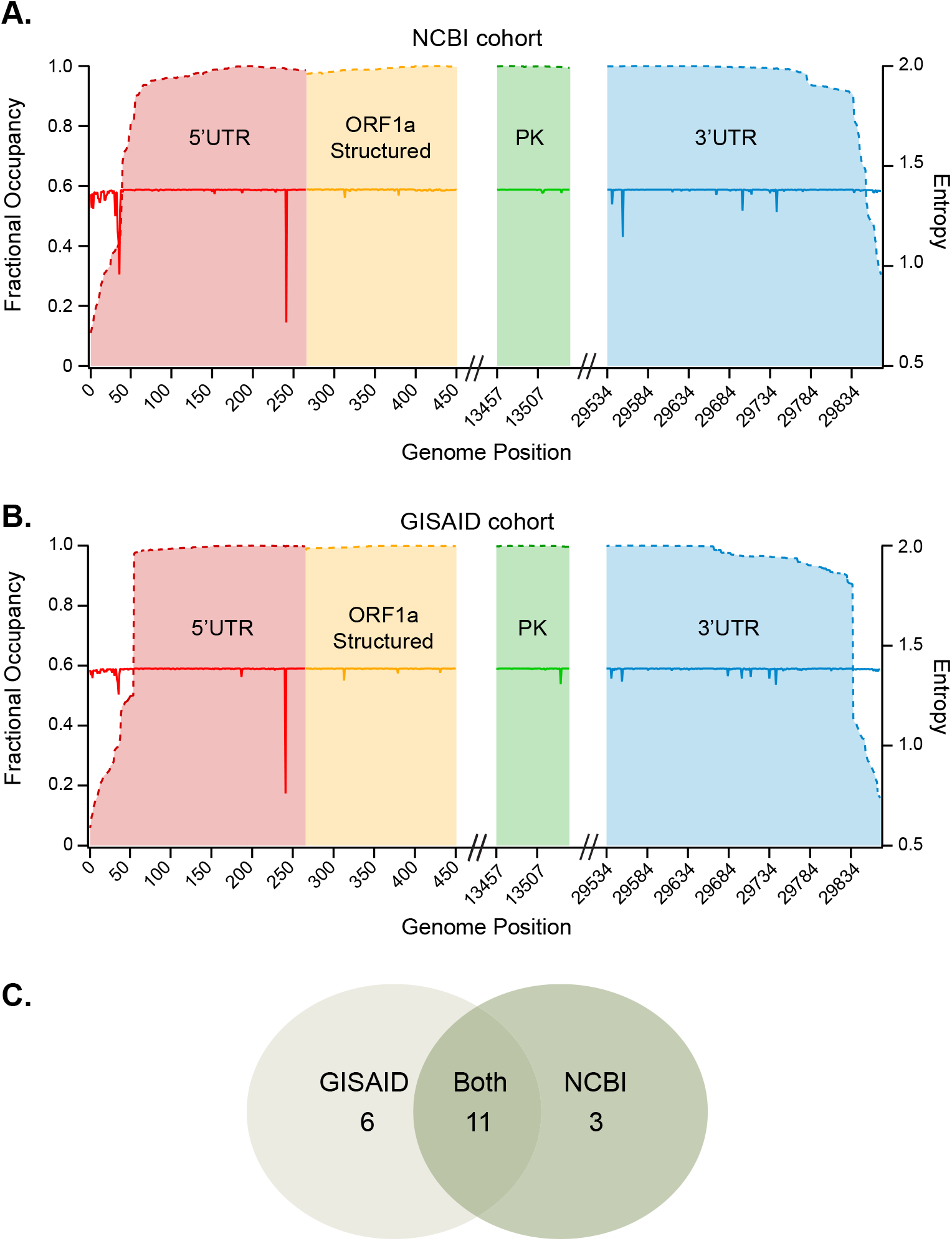
SARS-CoV-2 sequence occupancy and entropy. **A**. Fractional occupancy (left axis, dashed lines) and positional entropy (right axis, solid lines) of the NCBI cohort calculated by WebLogo3 as displayed as a function of SARS-CoV-2 genome coordinates. This analysis focused only on well characterized betacoronavirus structured elements. The relative positional relationships of each region are marked. Hash marks donate areas of the entire genome that were not considered in this study. **B.** The same representation as in (A), but calculated using the GISAID EpiCoV cohort. **C.** Venn diagram of rapidly emerging variations in the GISAID cohort, NCBI cohort, or both.

To extend this analysis, we repeated the study with a second cohort of SARS-CoV-2 sequences recovered from the GISAID database on May 13, 2020 (Elbe and Buckland-Merrett 2017; Shu and McCauley 2017). All sequences were downloaded from the database, converted into a blast library, then queried and analyzed as above with the NCBI cohort. An average of 23,900 ± 630 hits were recovered from each query. As with the NCBI cohort, occupancy is high (>90%) from position 55 through position 29,829 (Fig. 1B, Supplementary Table 1). Due to the large size of the GISAID cohort—6.6 times the size of the NCBI cohort—the termini are covered by thousands of genomes despite the relatively low occupancy. In total, seventeen variants with a MAF of at least 0.005 were identified in the GISAID cohort, eleven of which were also identified in the NCBI cohort (Fig. 1C, Supplementary Table 2). Combining the two analyses yields a total of twenty rapidly emerging variants in the structured regions of the viral genome. Of these, thirteen are transversions and seven are transitions. Eighteen are in noncoding regions

(2.9%, 18/602 positions evaluated), and the remaining two are silent mutations within the coding sequence of ORF1a or ORF1b (0.7%, 2/275 positions evaluated). Considering the larger GISAID cohort, there are 80 invariant residues (30.1%) in the 5’UTR, 90 (48.9%) in the ORF1a structured region, 58 (64.4%) in the frameshifting pseudoknot, and 131 (38.9%) in the 3’UTR. Thus, as expected, structures in the coding region seem to show a higher degree of conservation and fewer rapidly emerging alleles than noncoding regions, presumably due to the selective pressure of maintaining the protein coding sequence.

### Variations in SL1 through SL4 of the 5’UTR

In MHV, stem loop 1 (SL1) plays a critical role in virus replication and is proposed to form long-range interactions with the 3’UTR (Zuniga et al. 2004; Li et al. 2008). Stem Loop 2 (SL2) contains a highly conserved sequence and structural elements thought to play a role in sgRNA synthesis (Liu et al. 2007; Liu et al. 2009; Chen and Olsthoorn 2010; Lee et al. 2011). The structure of Stem loop 3 (SL3) is less well conserved, but it contains the leftmost transcription regulatory sequence (TRS-L) required for template switching in sgRNA production (Zuniga et al. 2004; Sola et al. 2005; Yang and Leibowitz 2015). Stem loop 4 (SL4) contains an upstream open reading frame (uORF) that could reduce translation initiation at the ORF1a start codon and/or act as a spacer between 5’ structured elements and ORF1a (Raman et al. 2003; Yang et al. 2011; Wu et al. 2014). The precise role of these structures in SARS-CoV-2 infection is not known, but recently released structural predictions reveal all four stem loops are present in the SARS-CoV-2 genome (Fig. 2) (Ramya Rangan 2020).

**Figure 2.**
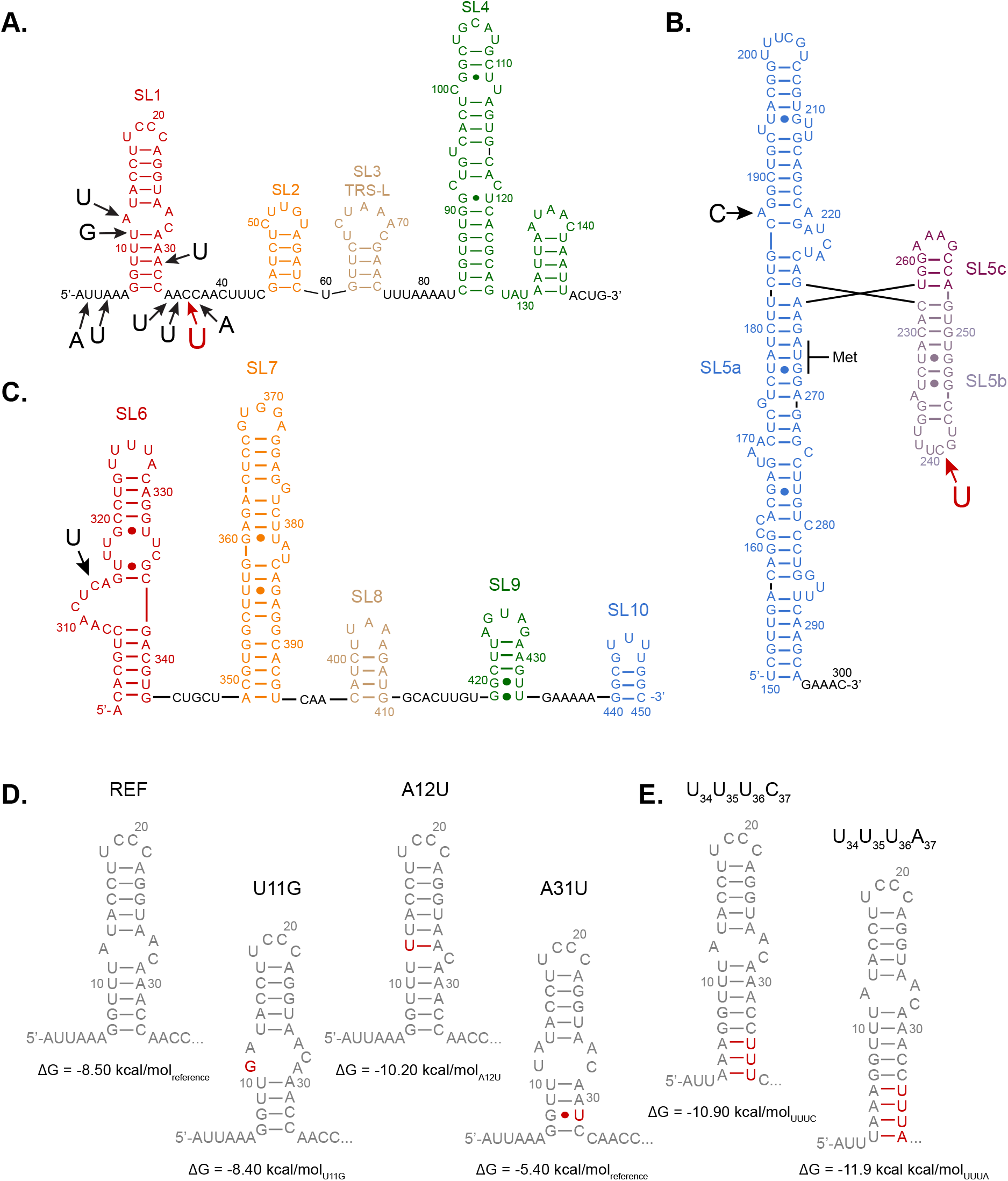
Rapidly emerging variants in SL1-SL10 of the 5’UTR and ORF1a: **A.** The predicted secondary structure of SL1 through SL4 is shown. The position and identity of rapidly emerging variants is denoted by an arrow and a letter. The minor allele frequency for each variant is given in the form variant=frequency/total (cohort). **B.** The predicted secondary structure of SL5 is shown. Rapidly emerging variants are denoted by an arrow, and the identity of the variation is given next to the arrow. The position of the ORF1a start codon is labeled. The minor allele frequency for each variation is given. **C.** The predicted secondary structure of SL6-SL10 is shown. Variations are labeled as in panel A. The minor allele frequency for the single variation is shown above, and includes the identity of the resultant silent codon change. **D.** The structures of single SL1 variants are shown. The specific variant is shown in red. The variant ID is given above the structure. The RNAfold calculated minimum free energy structure is presented in the diagram, and its thermodynamic stability is given below. **E.** Same as in D, but for the two prevalent combination variants that extend the length of SL1.

SL1 and flanking single-stranded regions contain nine of the twenty variants identified in this analysis. In contrast, no rapidly emerging variants (MAF >0.005) are found in SL2 through SL4. To determine how the variations in SL1 influence the secondary structure, we used RNAfold to calculate the most favored energy structure for each variant (Fig. 2D-E) (Lorenz et al. 2011). U2A (A=0.006/2519 GISAID, A=0.014/419 NCBI) and A4U (U=0.004/1799 GISAID, U=0.016/419 NCBI) have no influence on the stability of SL1 (ΔG=−8.50 kcal/mol for reference and both variants). U11G (G=0.001/4154 GISAID, G= 0.006/772 NCBI) had a small effect on the predicted stability (ΔG=−8.40 kcal/mol_U11G_) due to loss of a stem terminal A-U pair. The A12U variant (U=0.002/4484 GISAID, U=0.011/829 NCBI) stabilized the predicted structure through formation of an additional base pair (ΔG=−10.2 kcal/mol_A12U_). By contrast, A31U (U=0.005/7525 GISAID, U=0.029/1314 NCBI) is strongly destabilizing (ΔG=−5.40 kcal/mol), causing disruption of the lower stem.

Four rapidly emerging variations are found just downstream of the SL1 stem (Fig. 2A). A34U (U=0.009/7795 GISAID, U= 0.050/1350 NCBI), A35U (U=0.013/7846 GISAID, U=0.071/1380 NCBI), C36U (U=0.028/7967 GISAID, U=0.150/1400 NCBI) and C37A (A=0.003/8091 GISAID, A=0.018/1474 NCBI) variants frequently occur in combination. In the most common combination, three of the four positions (A34A35C36C37) are simultaneously replaced (U34U35U36C37). This variant has an allele frequency of UUUC=0.004/7795 (GISAID) and UUUC=0.023/1350 (NCBI). The variation extends the lower stem of SL1 by three base pairs, stabilizing the duplex by 2.4 kcal/mol (ΔG=−10.9 kcal/mol) (Fig. 2E). The second most frequent combination (U34U35U36A37) is present at an allele frequency of UUUA=0.003/7795 (GISAID) and UUUA=0.018/1350 (NCBI). This variant extends the SL1 lower stem by yet another base pair, increasing its overall stability by 3.4 kcal/mol (ΔG=−11.9 kcal/mol). Of these four positions, only C36U frequently exists as a single variation (U=0.013/7967 GISAID, 0.071/1400 NCBI). RNAfold analysis reveals no change in the stem loop structure or stability for the C36U variant.

Both combination variations are only found in samples sequenced from the United States, with the majority of them coming from the state of Washington. To better assess the relatedness between genomes containing U_34_U_35_U_36_C_37_ and U_34_U_35_U_36_A_37_ variants, we recovered the entire genomes of each example containing either extended SL1 stem variation from the GISAID cohort and aligned them using MAFFT (Katoh et al. 2002). The reference genome (Wuhan-Hu-1) was used as an outgroup. A radial maximal likelihood phylogenetic tree was calculated using the Tamura-Nei model in MEGAX, and the results plotted (Supp. Fig. 1) (Tamura and Nei 1993; Stecher et al. 2020). The phylogenetic relationship shows that both variants are represented in two different branches, but the variants tend to cluster separately within those branches. In one case, thirteen U_34_U_35_U_36_A_37_ genomes cluster within a node that is otherwise occupied U_34_U_35_U_36_C_37_, suggesting that U_34_U_35_U_36_ variation arose first, and A_37_ arose as a secondary mutation. The impact of these variations on viral fitness or patient outcomes is not known.

Most of the rapidly emerging variants in SL1 enhance the stem loop structure. This suggests that SL1 stabilization is not overly deleterious to virus replication. In MHV, by contrast, destabilizing mutations of the lower stem are well tolerated in a cell model of virus replication, but mutations that increase the stability the lower stem block replication (Li et al. 2008). It is important to note that there is significant sequence divergence in this region between the two viruses that may explain this apparent dichotomy. Interestingly, the combination U_34_U_35_U_36_C_37_ variation co-occurs with the destabilizing A31U mutation 48.5% of the time, suggesting a potential compensatory role. Consistent with this hypothesis, the extension in SL1 rescues the destabilizing A31U variation by 2.1 kcal/mol (ΔG=−7.5kcal/mol). However, we note that none of the combination U34U35U36A37 variant genomes harbor A31U, so it is clear that the SL1 extension can exist in the absence of a compensatory destabilizing mutation. There are no rapidly emerging variations within the upper stem or the loop of SL1, suggesting this region could be important to infection. Consistent with that hypothesis, mutations that destabilize the upper stem of MHV SL1 block virus replication (Li et al. 2008).

### Variations in SL5 through SL10 at the 5’UTR/ORF1a junction

A large branched helical structure termed Stem Loop 5 is predicted to form at the interface between the 5’UTR and the N-terminal region of ORF1a (Fig. 2B) (Ramya Rangan 2020). This region contains three stems (SL5a, SL5b, and SL5c) connected by a helical junction. There is considerable sequence divergence among the coronaviridae in this structure, but the overall fold is largely preserved (Chen and Olsthoorn 2010). In SARS-CoV-2, the SL5a stem occludes the initiation codon for ORF1a, suggesting this structure must open prior to translation initiation. However, the SL5a stem is essential for virus replication in a bovine coronavirus (BCoV) model (Brown et al. 2007). The role of SL5c is more controversial, with one study demonstrating that the stem is dispensable (Yang and Leibowitz 2015), while a previous study showed that it is required (Brown et al. 2007).

We observed two rapidly emerging variants within the SL5 structured region with a minor allele frequency of greater than 0.005. The first, A187C (C=0.007/23832 GISAID, C=0.003/3407 NCBI), occurs within a bulged nucleotide of SL5a and is therefore not expected to alter the structure. The second, C241U (U=0.682/23760 GISAID, U=0.616/3376 NCBI) is in SL5b loop and is the most abundant rapidly emerging variant by far. There are no rapidly emerging mutations with an allele frequency of >0.005 in SL5c in either cohort.

Four additional stem loop structures (SL6-SL10) have been proposed within ORF1a (Fig. 2C) (Ramya Rangan 2020). The presence of SL6 and SL7 is observed in other coronaviridae, but the structures do not appear to have an important function (Brown et al. 2007; Yang et al. 2015). There is one rapidly emerging variant within this region. C313U occurs within an internal loop region of SL6. The minor allele frequency of this variant is U=0.011/24227 GISAID, U=0.007/3732 NCBI. The variant is a silent mutation, converting a CUC^Leu^ codon to a CUU^Leu^ codon. As it occurs in an internal loop, it is expected to have no impact on the stem loop structure.

### Variations in the 5’UTR that could have arisen through RNA editing

The two most abundant variants in the 5’UTR are both C to U transitions. C36U is observed in 2.8% of the sequences from the GISAID cohort, and C241U is observed in 68%. Excluding singletons, the average frequency of C to U transitions at all other positions in the 5’UTR is 0.04%. It is possible that the C36U and C241U variations arise repeatedly during virus replication, or they may have occurred early during the outbreak, or both. The type of transition and the relative abundance of the C36U and C241U variations suggest they might be hot spots for viral genome editing by host defense enzymes. The apolipoprotein B mRNA editing enzyme catalytic polypeptide-like (APOBEC) enzymes are host encoded cytidine deaminases that edit cytidine to uridine in host nucleic acids (Lerner et al. 2018; Silvas and Schiffer 2019). They also target single stranded RNA and DNA virus genomes to affect an antiviral response.

If C36U and C241U substitutions arose at such high frequency because of C to U RNA editing, it might be possible to observe both nucleotides in the same sample of genomic RNA. cDNA produced from a mixed population of viral RNA harvested from an individual would be expected to include a weighted average of C and U in the sequencing reads that could be indicated as a degenerate Y (pyrimidine) in sequencing data, especially if there are near equal reads of each variation. Because WebLogo3 does not consider degenerate sequencing calls in its calculation of allele frequency (Crooks et al. 2004), we used SNP-sites v2.5.1 and VCFtools v0.1.7 to recalculate the allele frequency inclusive of degenerate bases (Danecek et al. 2011; Page et al. 2016). We calculated the average frequency of all C to Y transitions in the 5’UTR using the larger GISAID cohort, excluding the two candidate editing sites (C36U and C241U) and singletons. The average C to Y transition frequency is 0.014%. By contrast, the frequency of C36Y is 0.063%, 4.5-fold greater than the average, and the frequency of C241Y is 0.18%, 12.9-fold greater than the average. This apparent increase in pyrimidine degeneracy is consistent with the possibility that APOBEC enzymes edit both positions. However, we cannot formally rule out the possibility that some people were co-infected with both variants leading to the degenerate base call, or that C36U and C241U frequently arise via some other mechanism during viral replication. The impact of either variation on viral fitness remains to be determined.

### Variations in the frameshifting pseudoknot at the ORF1a/ORF1b junction

An RNA pseudoknot is found at the junction of ORF1a and ORF1b (Fig. 3A) (Ramya Rangan 2020). This structure is involved in −1 programmed ribosome frameshifting, where translating ribosomes shift frame by one nucleotide to the left. Efficient frameshifting requires both a “slippery” sequence and a downstream stable RNA structure (Brierley et al. 1989; Brierley et al. 1992). Like SARS-CoV and MHV, the SARS-CoV-2 pseudoknot has three stems instead of two typically found in pseudoknot structures (Brierley and Dos Ramos 2006; Giedroc and Cornish 2009). A previous study comparing SARS-CoV, MHV, and hybrid variants found that both viral pseudoknots led to approximately the same extent of programmed frameshifting (~20%), but hybrid mutant variants in loop 3 that stabilize the pseudoknot structure increased frame shifting up to 90% (Plant et al. 2010). The same study revealed that silent mutations in the SARS-CoV slippery site reduced programmed frame shifting by three-fold and also blocked viral infection in a cell culture model. Thus, the function of the slippery sequence and the pseudoknot structure is to ensure that production of ORF1a and ORF1ab polyproteins occurs at appropriate stoichiometric ratios, critical to viral fitness.

**Figure 3.**
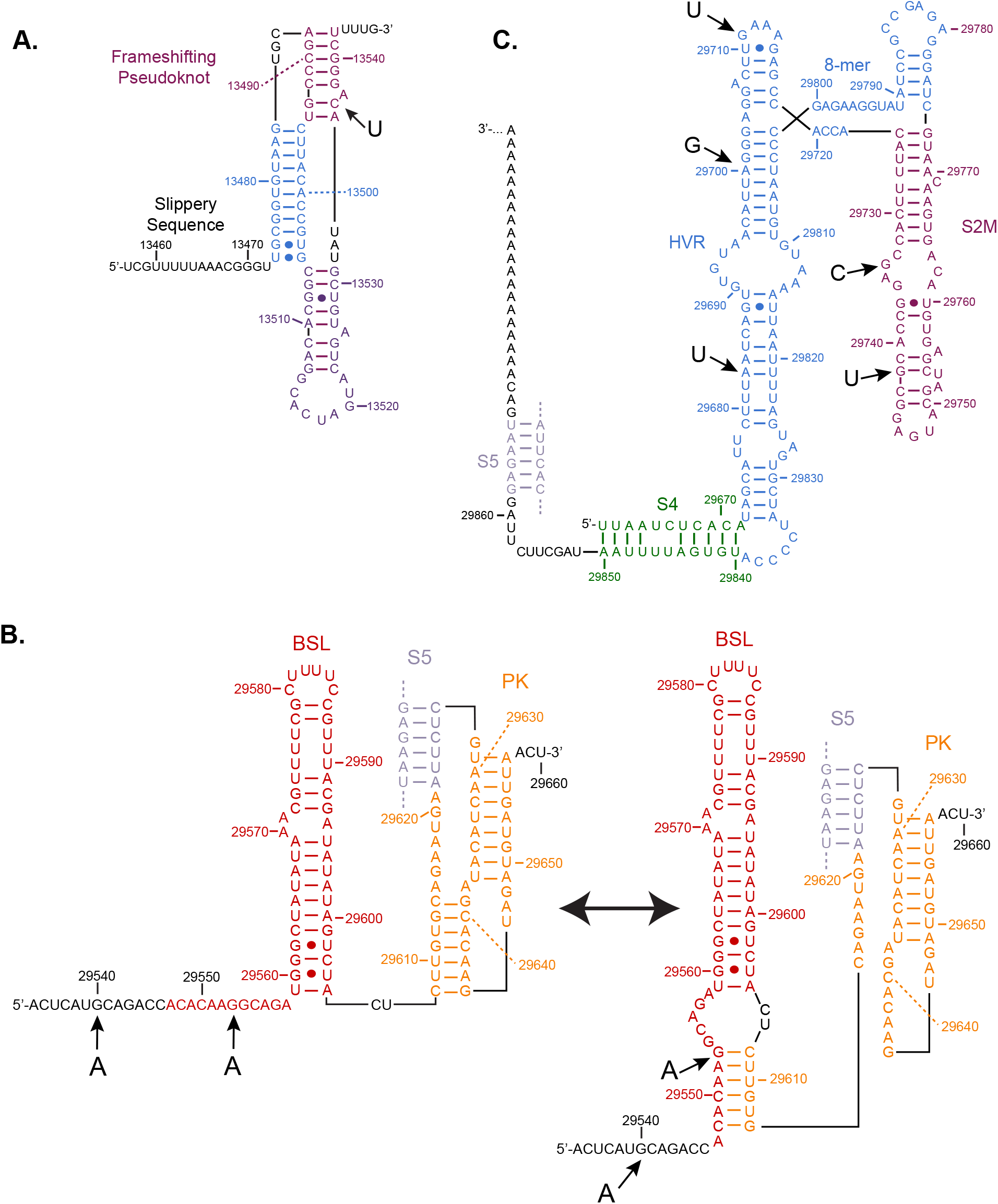
Rapidly emerging of the frameshifting pseudoknot and the 3’UTR: **A.** The secondary structure of the frameshifting pseudoknot is shown. **B.** The secondary structure of the pseudoknot conformer or the extended BSL conformer is shown of the bi-stable molecular switch in the 3’UTR is shown. **C.** The secondary structure of the 3’ half of the SARS-CoV-2 genome is shown. This region includes the HVR, the octanucleotide motif (8-mer), the S2M structure, and the S4 and S5 stems. In all paneles, the position of rapidly emerging variants is labeled as in figure 2.

We identified one rapidly emerging variant in the frameshifting pseudoknot. C13536U (U=0.015/23306 GISAID, U=0.003/3440 NCBI) is a silent mutation (UAC^Tyr^:UAU^Tyr^) located within stem 2 (Fig. 4). C13536 normally forms a Watson-Crick pair with G13493. Mutation to U is expected to cause the formation of a U13536:G13493 wobble pair, which has comparable stability to a Watson-Crick pair but alters the backbone geometry shifting the G residue into the minor groove. To get a better understanding of how this U-G pair might impact the tertiary structure and thus the function of the frameshifting pseudoknot, we used RNAcomposer to build a threedimensional model of the reference sequence and the C13536U variant (Fig 4B) (Popenda et al. 2012). In the reference model, G13485 forms a base triple with the C13536:G13493 pair (Fig. 4A). The exocyclic amine of C13536 donates a hydrogen bond to the O6 of G13485 in loop 1. In the C13536U model, this base triple cannot form as the hydrogen bond donor is lost. This could conceivably reduce the stability of stem 2, which would be expected to cause less efficient −1 programmed ribosomal frameshifting. More work will be necessary to define exactly how this variation perturbs the structure and if it alters the stoichiometry of viral protein synthesis.

**Figure 4.**
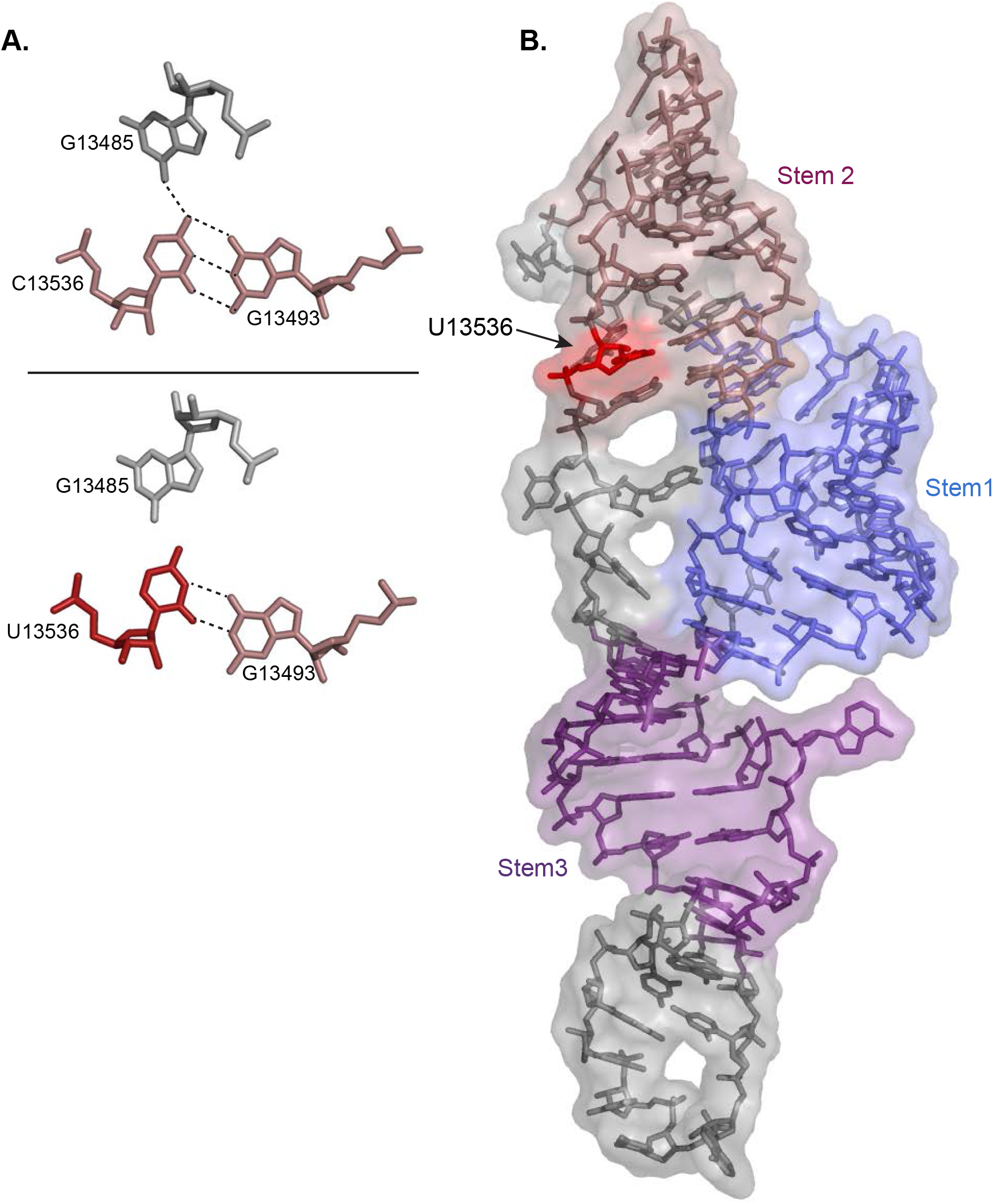
Structure of the frameshifting pseudoknot. **A.** Comparison of the base triple observed in the reference model (top) and in the U13536 variation model (bottom). Hydrogen bonds are denoted by dashed lines. The U13536 variant is colored in red. **B.** The molecular model of the frame shifting pseudoknot calculated by RNAcomposer is shown. Stems 1-3 are labeled in colors corresponding to those shown on the secondary structure in figure 3.

### Variations in the 3’UTR in the BSL and PK

Betacoronaviridae 3’UTRs contain a bi-stable molecular switch formed by two mutually exclusive structural conformers, including one that extends the lower stem of the bulged stem loop (BSL), and a second that folds into a pseudoknot (PK, Fig. 3B) (Goebel et al. 2004). Both structured elements are present in MHV, SARS, and MERS, though the sequence diverges significantly between them (Hsue and Masters 1997; Hsue et al. 2000; Goebel et al. 2004). In MHV and BCoV, the BSL and the PK structure are required for viral replication (Hsue and Masters 1997; Williams et al. 1999; Hsue et al. 2000; Goebel et al. 2004). Mutations that stabilize one form over the other prevent replication. It is proposed that competition between the two structures plays a regulatory role in antigenomic RNA synthesis, but the exact mechanism remains to be determined (Goebel et al. 2004).

There are two rapidly emerging variants in the 5’ portion of the 3’UTR. G29540A is present at a MAF of A=0.008/24313 (GISAID) and A=0.014/3550 (NCBI). This variant lies within a single-stranded region that precedes the BSL structure and as such is not predicted to affect the structure or the molecular switch. In contrast, the G29553A variant (A=0.012/24216 GISAID, A=0.064/3551 NCBI) disrupts a G:C pair in the extended BSL molecular switch conformer that could potentially favor the alternate PK structure. Alternatively, the A substitution may pair with the otherwise bulged U29607 nucleotide, partially compensating for the loss in of the G:C pair. Consistent with the latter possibility, RNAfold predicts that the stability of the reference BSL conformer is −20.20 kcal/mol, while the stability of the G29553A variant conformer is −18.30 kcal/mol and includes a newly formed A:U pair. It remains to be determined how modulation of the internal equilibrium of the molecular switch affects SARS-CoV-2 pathogenesis.

RNA editing by APOBEC enzymes could lead to rapidly emerging G to A transitions if the antigenomic strand is edited during viral replication. Antigenomic cytidine deamination recodes C to U, which would be read as an A during replication of the genomic strand. To assess this possibility that the G29540A and G29553A variations arose through RNA editing, we looked for the degenerate base “R” (either purine base) in both sequencing cohorts using SNP-sites and VCFtools as described above (Danecek et al. 2011; Page et al. 2016). There were no degenerate R alleles in the GISAID or NCBI databases at either position, suggesting that neither is produced through frequent APOBEC-mediated editing of the antigenomic strand.

### Variants of the hypervariable region, the S2M structure, and the S3 and S4 stems

An extended multiple stem loop structure exists downstream of the 3’UTR pseudoknot (Fig 3C) (Ramya Rangan 2020). This structure contains a hypervariable region (HVR) that folds into a bulged stem loop. The HVR is highly divergent in coronaviridae with the exception of a strictly conserved single-stranded 8-mer sequence referred to as the octanucleotide motif (Goebel et al. 2007; Madhugiri et al. 2014). The function of this region is not well understood, but deletion of the HVR including the conserved 8mer element has no effect on MHV replication in cultured cells (Goebel et al. 2007). An apparent selfish genetic element, termed S2M, exists within the bulged stem loop of the HVR (Ramya Rangan 2020). This element is found in many but not all coronaviridae, and is also found in many other families of positive ssRNA viruses, suggesting it can be horizontally transferred (Tengs et al. 2013; Tengs and Jonassen 2016). The sequence is highly conserved in all viruses where it is found. This element is not present in MHV, and its function (if any) is unknown. Two shorter stems, termed S4 and S3, are also present. Mutations that disrupt S4 have no effect on MHV replication, but S3 appears to be important (Liu et al. 2013).

Five rapidly emerging variants are found in this region of the SARS-CoV-2 genome. Two disrupt Watson-Crick pairs in the HVR bulged stem loop. The A29683U variation is present at a MAF of U=0.006/23540 (GISAID) and U=3×10^−4^/3545 (NCBI), while the A29700G is present at a MAF of G=0.009/23403 (GISAID) and G=0.022/3541 (NCBI). Both variants reduce the stability of a simplified model HVR structure that eliminates the S2M region in RNAfold calculations (ΔG=−24.20 kcal/mol_ref_, –22.20 kcal/mol_A29683U_, –23.9 kcal/mol_A29700G_, Supplemental Fig. 2), with the A29700U variant forming a compensatory G:U wobble pair. The G29711U variant is present at a MAF of U=0.007/23366 (GISAID), U=0.004/3537 (NCBI). This variant disrupts the GNRA class tetraloop structure in the loop of the HVR bulged stem structure, and is predicted to modestly destabilize the fold (ΔG = −23.5 kcal/mol_G29711U_). The presence of multiple disruptive variations in this region of the SARS-CoV-2 3’UTR, coupled to previous reports that the HVR is dispensable for MHV replication, suggests that structures are not critical to viral replication. More work will be needed to understand whether the structures in the HVR contribute to SARS-CoV-2 replication or viral fitness.

The presence of the rapidly emerging A29700G transition suggests the possibility that it might arise through adenosine deaminase acting on RNA (ADAR) RNA editing activity. ADARs convert adenosine residues to inosine in double stranded regions of RNA (Keegan et al. 2017). As such, they can play an important role in antiviral response, targeting double stranded RNA viruses and other viruses (including betacoronaviruses) that go through a double stranded RNA intermediate (Tomaselli et al. 2015). During viral replication, inosine residues in the genomic strand would template the incorporation of a C in place of a U during minus strand synthesis, leading to A to G transitions during viral replication. As above, we used SNP-sites and VCFtools to measure the frequency of the degenerate R base at A29700G (Danecek et al. 2011; Page et al. 2016). No degenerate R nucleotides are present in the GISAID cohort, suggesting that frequent RNA editing by ADAR enzymes is not responsible for rapid A29700G emergence.

The final two rapidly emerging variations lie within the enigmatic S2M loop. The structure of the S2M loop from SARS-CoV has been solved by X-ray crystallography (Fig. 5) (Robertson et al. 2005). Its prevalence in positive strand ssRNA viral genomes, its position near the 3’-terminus, and its high degree of sequence conservation all imply a functional role (Tengs and Jonassen 2016). However, not all betacoronaviruses have the S2M loop, and swapping an S2M-containing region from the SARS-CoV 3’UTR with an S2M-deficient MHV region did not alter or improve virus replication in vitro (Goebel et al. 2007). As such, its role in viral replication is unclear.

**Figure 5.**
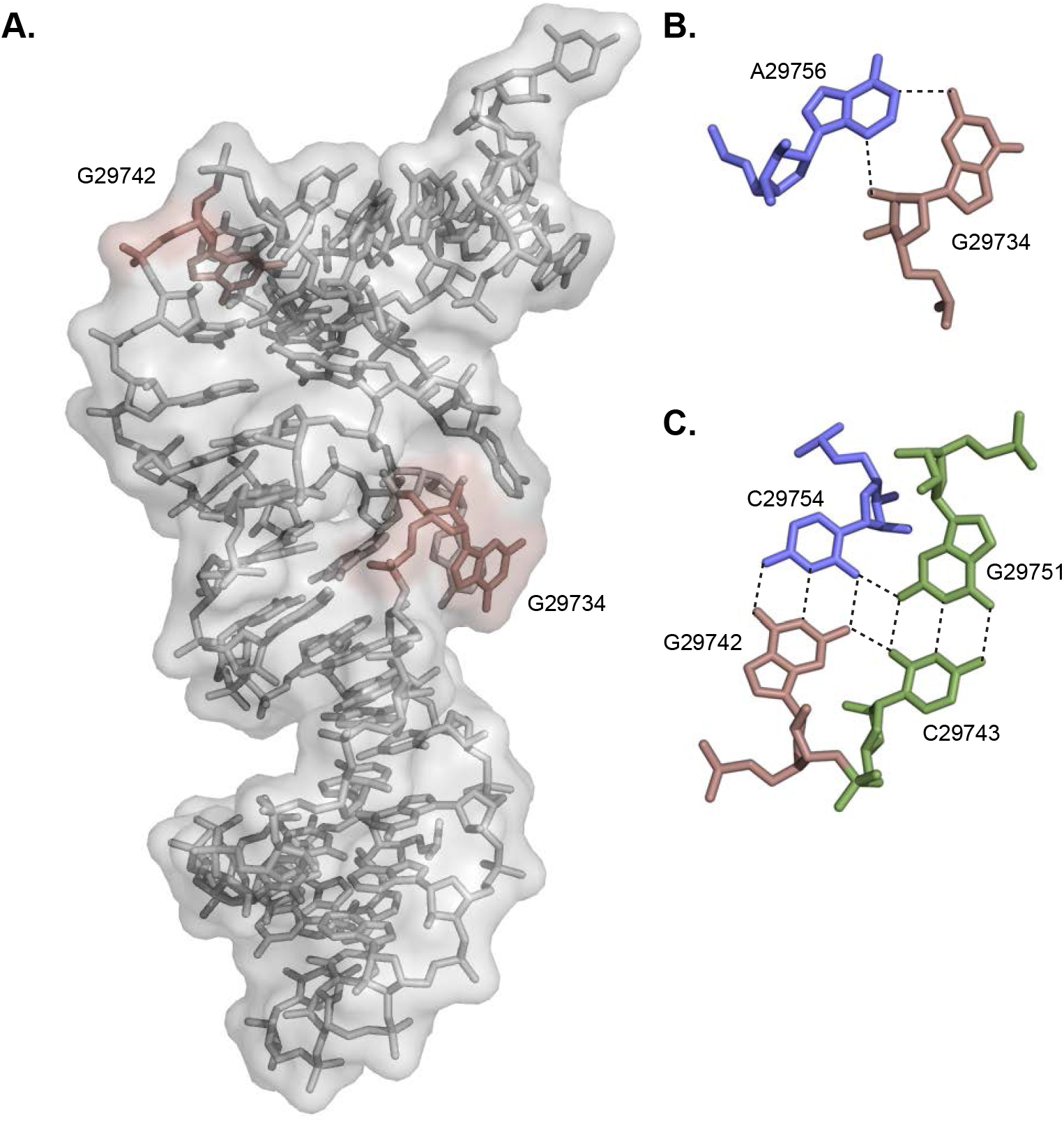
Rapidly emerging variations in S2M disrupt key tertiary interactions: **A.** The crystal structure of the S2M region from SARS-CoV is shown. The position of two rapidly emerging variations in SARS-CoV-2 are shown in red adjacent to corresponding nucleotides in the SARS-CoV structure. **B.** G29734 is involved in a non-Watson-Crick pair with A29756. The hydrogen bonding pattern is denoted with dashed lines. Both nucleotides are conserved between SARS-CoV and SARS-CoV-2. The variant position is marked with red **C.** G29742 is involved in a base quadruple with three residues conserved between SARS-CoV and SARS-CoV-2. The Watson-Crick partner of G29743 is in blue. A G:C pair that packs into the minor groove is shown in green. The position of the variant base is denoted by red. Hydrogen bonds between the bases are shown as dashed lines.

The first rapidly emerging variation in S2M is G29734C (C=0.008/23285 GISAID, C=0.003/3525 NCBI). In the SARS-CoV S2M crystal structure, this position forms a non-canonical G:A pair (Fig. 5B) (Robertson et al. 2005). The N2 exocyclic amine donates a hydrogen bond to the N1 position of its adenosine partner, and the 2’-hydroxyl group donates a hydrogen bond to the N3 moiety. Substitution of a C in place of G is incompatible with the hydrogen bonds formed in the G:A pair and as such is likely to destabilize the S2M tertiary structure. The second rapidly emerging variation in S2M is G29742U (U=0.009/28235 GISAID, U=0.019/3526 NCBI). This base is involved in a base quadruple, pairing through its Watson-Crick face with a cytidine residue, but also interacting with the C of a parallel G:C pair packed tightly into its minor groove (Fig. 5B) (Robertson et al. 2005). The U variation is incompatible with both the canonical and non-canonical pairings at this position and is likely to be highly destabilizing to the fold.

To evaluate the effect of both of the rapidly emerging variations on the structure of the S2M loop, we performed molecular dynamics (MD) simulations of the S2M loop from SARS-CoV and of both variants. The X-ray structure of the SARS-CoV S2M loop served as initial conformation for all simulations (Robertson et al. 2005). The results confirmed our predictions: both variations destabilize the structure of the S2M domain. In each case we observed that the structure of the S2M loop undergoes a transition quickly after equilibration (within the first 60 ns) to a less compact conformation as shown in Figure 6A-F. To quantify the extent of the transition, we measured the length of the maximum dimension of the S2M loop and observed an extension (Fig. 6I-K). Surprisingly, we observed that the S2M loop of SARS-CoV also samples similarly extended conformations (Fig. 6I), although this happens more slowly and with a lower frequency than in the other two variants, in only two out of the four trajectories that we collected (Supplemental Fig. 3A-C). Analysis of the MD trajectories showed that one important element for the stability of the overall structure of the S2M loop is the stabilizing interaction between G29734 and A29756. When these two nucleotides are in close proximity to one another, the stem tertiary structure is reorganized by bending. This interaction is destabilized in the G29734C variant because C cannot form hydrogen bonds with A29756. However, we observed that the formation of an A-G base pair is not stable in all three sequence variations (Fig. 6G). In the absence of this base pair, A29756 can move far from G/C29734 (Fig. 6L-N) leading to the sampling of more extended conformations of the loop as shown in figure Fig6 C-F. With the exception of the tails that unfold and refold, the secondary structure of the S2M loop is generally well preserved throughout the simulations, even when sampling more extended conformations as a result of the loss of tertiary interactions (Fig. 6G). A major difference between these variants is that the base pairing is not as stable in the stem-loop structure in the presence of the G29742U variation (Fig. 6G-H). This variation in sequence affects the stability of the base quartet described above (Fig. 5A), and ultimately impacts the stability of the adjacent base pairs (Fig. 6H). The base quartet is at the junction between the two helices and is important to set their relative orientations. The G29742U variation causes a local melting of the secondary structure and a disruption of this junction, ultimately destabilizing the compact conformation of the S2M loop (Fig. 6C).

**Figure 6.**
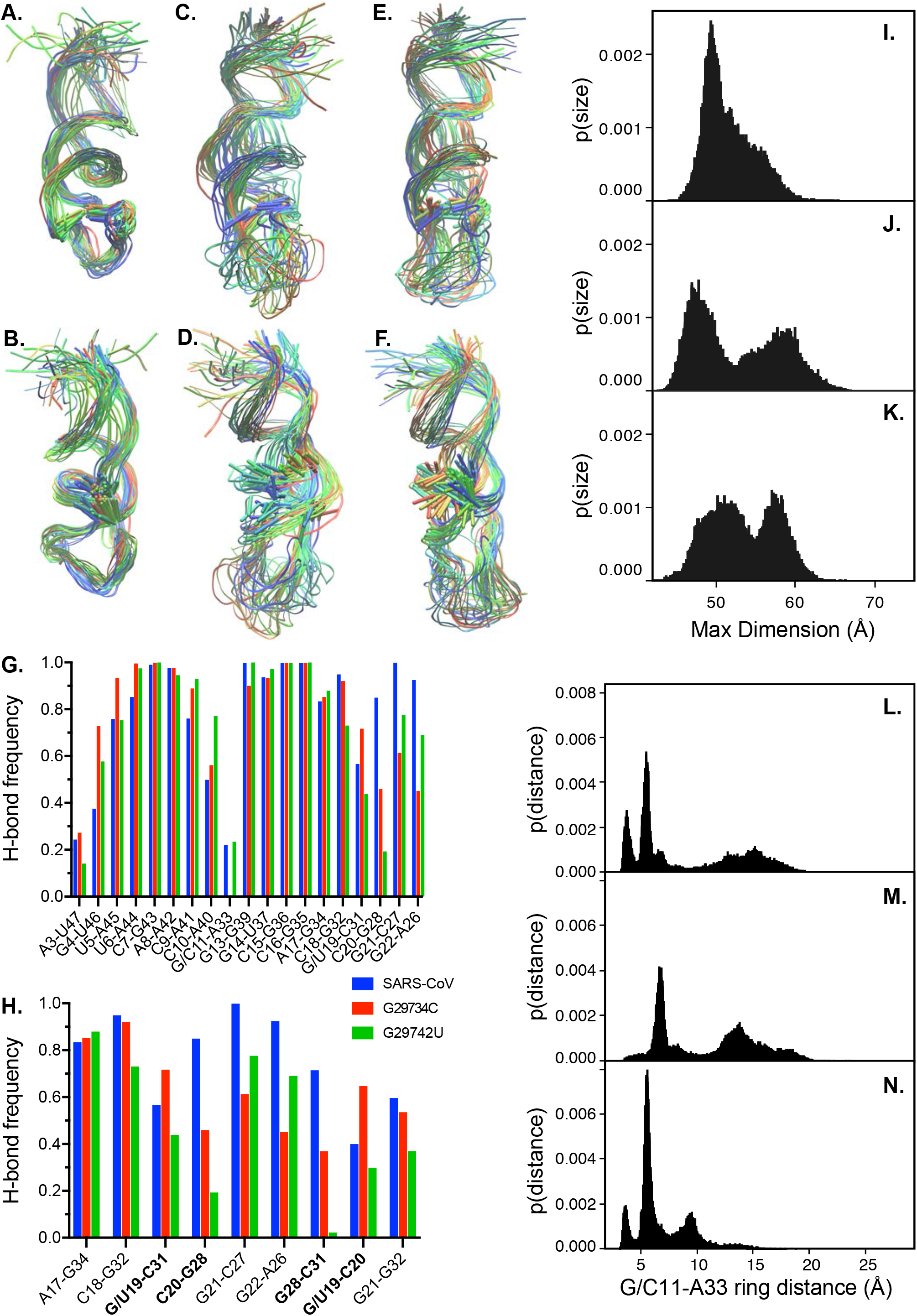
Molecular Dynamics simulations of S2M variations. Overlay of the structures from a single representative 180 ns trajectory of the SARS-CoV S2M loop (**A-B**), G29734C (**C-D**) and G29742U (**E-F**) variants. Front and back orientations show the following residues as sticks: G/U29742 and C29754 in A, C and D; G/C29734 and A29756 in B, D and F. Structures are colored as a function of time (blue = 0 ns, red =180 ns). Hydrogen bond frequency between the base pairs of the S2M loop is shown in **G** for SARS-CoV (blue), G29734C (red) and G29742U (green) variants. Hydrogen bond frequency for the interacting nucleotides in the quartet (highlighted in bold font) and base pairs around G/U29742 is shown for SARS-CoV (blue), G29734C (red) and G29742U (green) variants in **H**. The hydrogen bond frequency is calculated over the four 180 ns trajectories in G and H. Histograms of the largest dimension of the S2M loop measured for the four 180 ns trajectories of SARS-CoV (**I**), G29734C (**J**) and G29742U (**K**) variants. Histograms of the base distance measured between G/C29734 (G/C11) and A29756 (A33) for the four 180 ns trajectories of SARS-CoV (**L**), G29734C (**M**) and G29742U (**N**) variants. In all panels the nucleotides are numbered as in the X-ray structure of SARS CoV S2M RNA (PDB code 1XJR), the corresponding number in the reference genome can be obtained by adding 29,723.

## Discussion

The global SARS-CoV-2 pandemic has led to an explosion in whole genome sequencing of naturally occurring viral isolates. These data have been useful in the identification of rapidly emerging variations that impact viral protein structure and function (Kim et al. 2020b; Pachetti et al. 2020). They have also been used to monitor the spread of the virus through molecular phylogeny (Chu et al. 2020; Coronaviridae Study Group of the International Committee on Taxonomy of 2020; Forster et al. 2020).

Here, we have used available data to investigate how rapidly emerging variants could impact structured cis-regulatory elements in the virus genome. These elements govern viral replication, subgenomic RNA synthesis, and translation control in other betacoronaviruses (Yang and Leibowitz 2015; Madhugiri et al. 2016). Rapidly emerging variants could enhance or dampen viral pathogenesis and overall fitness, which could affect the extent and duration of the outbreak. As such, it is critically important to understand how such variations arise, and what regions of the genome are most prone to mutation.

Due to the burden of the SARS-CoV-2 outbreak, there is renewed interest in the development of novel strategies to treat betacoronavirus infections. Functional RNA structures in the viral genome could provide new targets for small molecule therapeutic development. Many antibiotics work through interactions with ribosomal RNA structure, and RNA targeting small molecule drugs are currently approved or in development for a variety of infectious and genetic diseases (Guan and Disney 2012). The SARS-CoV-2 genome has many structured elements that could be targeted, including SL1-SL4 in the 5’UTR, the frameshifting pseudoknot at the ORF1a and ORF1b boundary, and the molecular switch in the 3’UTR. The results presented here suggest that the hypervariable region, including the S2M structure, might be less well suited to targeted drug development. Structures with rapidly emerging variations are problematic for drug development as well, as the relatively high viral mutation rate, coupled to its potential to be edited by APOBEC and ADAR enzymes, could lead to the rapid evolution of resistant variants.

Similarly, hybridization-guided therapeutics, such as antisense oligonucleotides, small interfering RNAs, and CRISPR-derived drugs could potentially be targeted to the SARS-CoV-2 genome. Unstructured regions in noncoding regions of the viral genome make particularly compelling targets, as access will not be blocked by RNA structure or transit of the ribosome. However, because these strategies rely on base complementarity to achieve target specificity, rapid virus evolution could prove their Achilles’ heel. The data presented here identify regions less prone to variation, making them better candidates for RNA-guided therapeutics.

The observation that SL1 is prone to rapidly emerging variations is interesting, as this region is not only present on the positive strand of the viral genome, but is also found on all subgenomic RNAs (Kim et al. 2020a). Moreover, the complement to SL1 in antigenomic RNA is likely recognized by viral RNA-dependent RNA polymerase (RdRP) to produce genomic copies of the viral RNA. As such, it could make a good target for therapeutic development. However, the presence of multiple variations, often in combination, makes strategies that rely upon base pairing unlikely to be effective for all virus subtypes. The diversity of variations that enhance the stability of SL1, including variations that lengthen the stem, suggests that SL1 stability is important to SARS-CoV-2 replication. But if stability matters more than sequence identity, we can expect the evolution of rapid resistance to therapeutics designed to modulate SL1 stability.

The bi-stable molecular switch in the 3’UTR is potentially the most compelling structure for targeted drug discovery. It is conceptually straightforward to design antisense oligonucleotides that lock the switch into one conformer or the other. Both conformers are necessary for MHV replication, and only one rapidly emerging variant of minimal consequence was identified in this region. It is likely that this switch plays a role in SARS-CoV-2 replication, as has been observed in other betacoronaviruses. More work will be necessary to assess its potential as a drug target.

RNA editing appears to play a role in two rapidly emerging variations near stem loop structures in the 5’UTR. The prevalence of RNA editing of the viral genome is not known, and it remains unclear whether editing affects viral fitness or pathogenesis. It will be interesting to assess the extent of RNA editing during active infection, a task that would probably be best achieved through direct RNA sequencing (Kim et al. 2020a).

The analyses presented in this study will only improve as more sequencing data are added to available repositories (Elbe and Buckland-Merrett 2017; Shu and McCauley 2017). It is possible that identification of more rapidly emerging variants will clarify some of the remaining ambiguities. The results presented here highlight the power of high-throughput sequencing of viral genomes to define viral cis-regulatory elements, and stand as a testament to the researchers collecting, sequencing, and sharing viral genomic data to help quell the impact of this tragic and overwhelming pandemic.

## Materials and Methods

### Calculation of allele frequency and occupancy

The Wuhan-Hu-1 isolate of SARS-CoV-2 (GenBank Accession Number MN908947) was used as a reference genome. The sequences corresponding to the 5’UTR (1-265), the ORF1a structured region (266-450), the frameshifting pseudoknot (13457-13546), and the 3’UTR (29534-29870) were used as queries in a BLASTN search (Altschul et al. 1990). For the NCBI cohort, BLASTN searches were performed against the <u>NCBI betacoronavirus database</u> of 11,495 (as of May 14^th^, 2020) betacoronavirus sequences. Searches were performed using the web portal with default parameters except “max target sequences” was set to 20,000. BLAST hits were filtered by organism for “severe acute respiratory syndrome coronavirus 2”, and the remaining hits were downloaded as a hit table and aligned sequences. A multiple sequence alignment was prepared using a locally installed copy of MAFFT version 7.464 using the default FFT-NS-2 algorithm (Katoh et al. 2002). The output file was then analyzed with a locally installed copy of WebLogo3 version 3.6.0 (Crooks et al. 2004). The resultant logo data table contains the calculated sequence entropy, the occupancy (weight), and the count number for each base at each position. The allele frequency was then calculated by dividing the count number by the sum of all counts for all four bases. The minor allele frequency is defined as the frequency of the second most abundant allele and is typically represented by the format variant=frequency/counts.

For the GISAID cohort, 24,468 curated SARS-CoV-2 genomic sequences were downloaded from the <u>GISAID Initiative</u> EpiCoV database (on May 13^th^, 2020) under the terms of their data access agreement (Elbe and Buckland-Merrett 2017; Shu and McCauley 2017). The genomic sequences were compiled into a blast library using a locally installed copy of BLAST+ version 2.8.1, and queried using the command line tool blastn as describe above with the exception that the max_target_seqs flag was set to 30,000 (Camacho et al. 2009). Aligned sequences were recovered from the resulting hit table using a custom shell command, then analyzed using MAFFT and WebLogo3 as described for the NCBI cohort above.

### Calculation of minimum free energy structures

The sequence corresponding to SL1 and flanking nucleotides (1-37), the BSL and flanking nucleotides (29,547-29,643), or variations thereof were input into the web server for <u>RNAfold</u> using the default parameters (Lorenz et al. 2011). The calculated ΔG for the minimum energy structure, the ensemble free energy, the frequency of the minimum free energy structure in the ensemble, the ensemble diversity, and the secondary structure in dot-bracket notation were recorded in Supplementary Table 3. The bulged stem loop in the HVR (29,627-29,834) and variants thereof were analyzed by the same approach, except nucleotides 29,721 through 29,800 were removed to simplify the overall structure. RNAfold was not able to accurately calculate the secondary structure of the region surrounding the s2m structure.

### Phylogenetic analysis of SL1 variants

Examples of the specific combination variants U34U35U36C37 and U34U35U36A37 were recovered from the GISAID cohort 5’UTR BLASTn hits by searching for the variation plus two invariant nucleotides on either side using custom shell commands. Each variant combination was searched using this approach to count the number of occurrences and to recover the sequence. Following alignment, the hits were inspected to ensure the correct pattern match, and in one instance, manually edited to remove an example where the search pattern identified a match at the incorrect position. The sequence IDs were then used to recover the intact genomic sequence from the GISAID cohort library. MAFFT was then used to generate multiple sequence alignments of the entire genome using the procedure outlined above. Output files were loaded into MEGAX version 10.1.8 (for Mac), and the maximum likelihood tree was calculated using the Tamura-Nei model (Tamura and Nei 1993; Stecher et al. 2020).

### Degenerate base frequency analysis

Because WebLogo3 does not consider degenerate base calls, the MAFFT-generated MSA files outlined above were converted into VCF format using a locally installed copy of SNP-sites version 2.5.1. The allele frequencies were then re-analyzed using VCFtools version 0.1.17 (Danecek et al. 2011; Page et al. 2016). The abundance of Y or R degenerate base calls for specific positions was calculated from the overall frequency each base, excluding counts for symbols that denote the absence of a base at the given position.

### Molecular modeling of the frameshifting pseudoknot and variants

Three-dimensional molecular models of the frame shifting pseudoknot (13,472-13,543) and variants thereof were calculated using the <u>RNAcomposer web server</u>. The modeling algorithm was guided using dot-bracket notation to match the recently published secondary structure of SARS-CoV-2 (Popenda et al. 2012; Ramya Rangan 2020). The output PDB files were visualized and analyzed in Pymol version 1.7.6.0.

### Molecular Dynamics simulations

Molecular dynamics simulations were performed with NAMD 2.13 (Phillips et al. 2005) using the CHARMM 36 force field (MacKerell et al. 1998). The X-ray structure of SARS-CoV S2M RNA (PDB code 1XJR) was used as the initial structure for the SARS-CoV and the G29734C and G29742U variants after performing the respective mutations using the Mutator plug-in of VMD (Humphrey et al. 1996). In VMD, each structure was solvated with a water box with explicit TIP3P (Jorgensen et al. 1983; MacKerell et al. 1998) solvent and an ionic concentration of 0.15 M NaCl. The box was much larger than the initial structure (cubic box of 106 Å) to allow for extension of the RNA. Four independent trajectories of each system (WT, G11C, and G19U) were generated with the following procedure. The solvated structures were first minimized using the conjugate gradient method for 500 steps to relax any high energy contacts or unphysical geometries. An additional 2,000 steps of conjugate gradient minimization were performed with the heavy atoms of the RNA and the two Mg^2+^ ions restrained with a harmonic constraint force of 10 kcal/mol^−1^ Å^2^. Next, particle velocities were randomly assigned from the Maxwell distribution and equilibration was performed in the isothermal-isobaric ensemble gradually decreasing the restraints to zero (using 9 stages of 50 ps each). The pressure and temperature were maintained at 1 atm and 298 K using Langevin dynamics and the Nosé-Hoover Langevin piston method. The SHAKE constraint algorithm (Ryckaert et al. 1977) was used to allow a 2 fs time step. The particle mesh Ewald method (Essmann et al. 1995) was used to calculate electrostatic interactions with periodic boundary conditions. Production trajectories were then collected using the isothermal-isobaric ensemble for 180 ns. The Mg^2+^ ions remained stably coordinated throughout the trajectories.

Trajectory analysis was performed with VMD (Humphrey et al. 1996) and structures were visualized with VMD using the STRIDE algorithm for secondary structure identification (Frishman and Argos 1995). A hydrogen bond is defined by a donor-acceptor distance of less than 3.5 Å and a donor-hydrogen-acceptor angle of 130° < θ < 180°. The ring distance between a pair of nucleotides was calculated as the center of mass distance between the atoms N1, C2, N3, C4, C5, and C6. The maximum dimension of the RNA structure was calculated by aligning the trajectories with the starting structure (where the longest axis of the RNA was aligned with the y-axis) and finding the maximum dimension of the box needed to fully contain the RNA.

## Acknowledgements

We thank and gratefully acknowledge the originating laboratories responsible for obtaining specimens and the submitting laboratories that generated genetic sequencing data shared through the GISAID Initiative. A full list of these laboratories is presented in the supplementary table 4. We thank Jeremy Luban, Brian Kelch, and members of the Ryder laboratory for commenting on the manuscript and for helpful discussions. This work was supported in part by the COVID19 Pandemic Research Fund at UMass Medical School. Research in our labs is funded by NIH grant R21HG011001 and R01GM117237 to S.P.R. and NIH grant R01 GM139316 to F.M. and S.P.R.

## Author Contributions

SPR performed the sequence analyses and molecular modeling. BRM collected the MD trajectories. BRM and FM performed the MD trajectory analyses. SPR wrote the paper under the advisement of all authors.

## Supplemental Figure Legends

**Supp. Figure 1.**
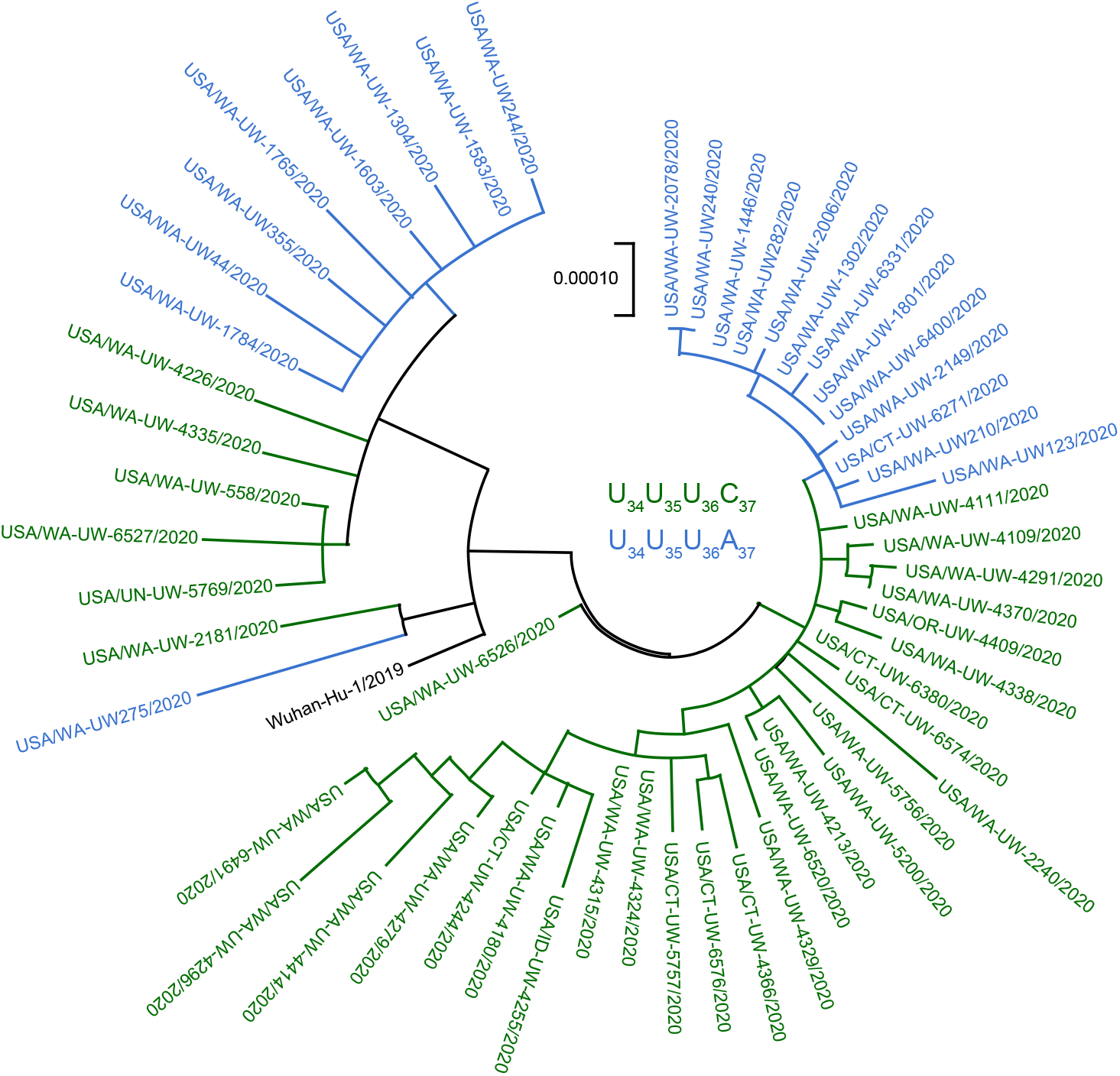
Maximal Likelihood phylogenetic tree calculated using the entire genome sequence of isolates carrying the SL1 variations in Fig. 1E. UUUA variants are colored blue, and UUUC variants are colored green.

**Supp. Figure 2.**
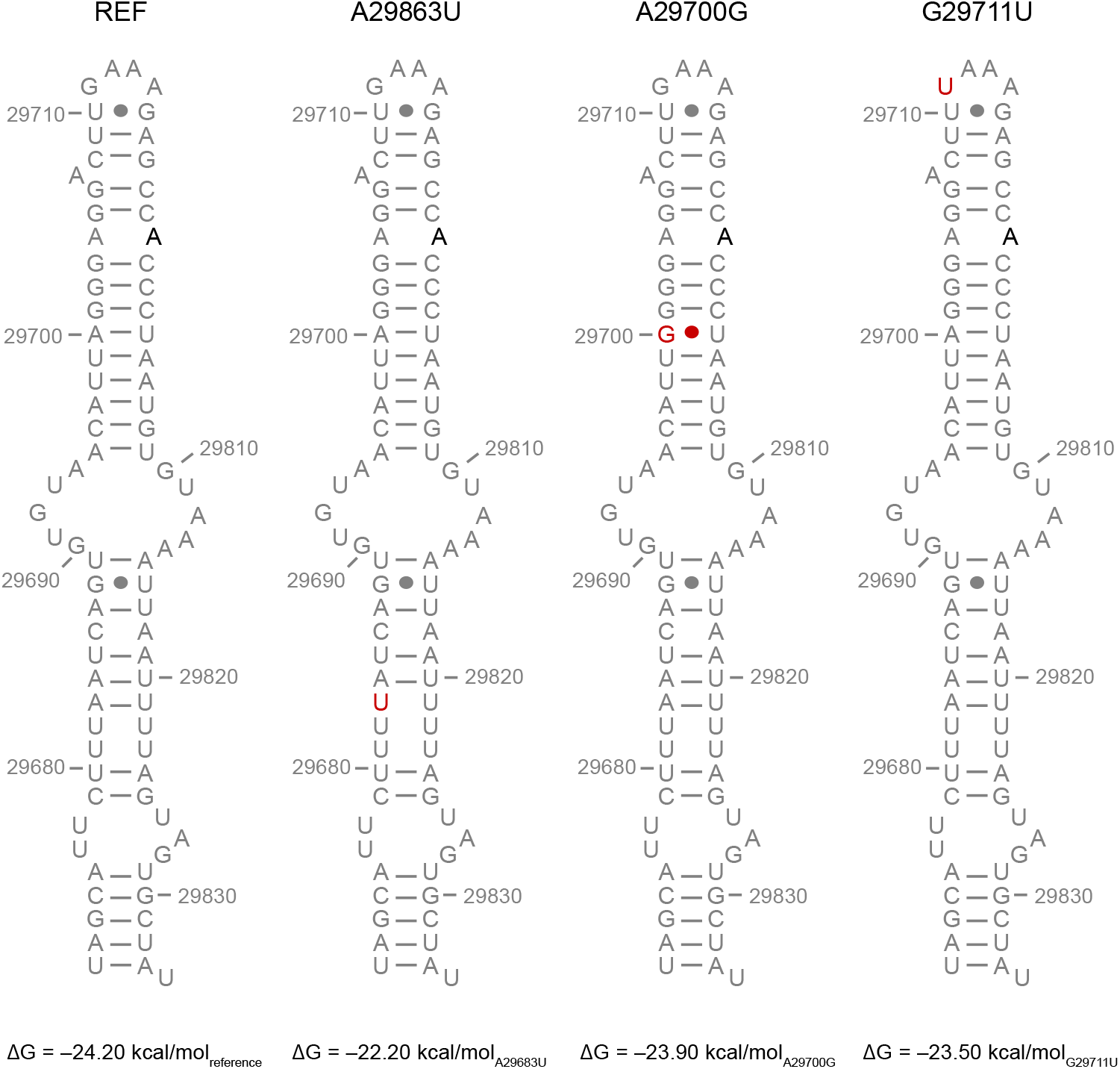
Most HVR variations destabilize the bulged stem loop structure. The secondary structure of a simplified HVR stem is shown. The region containing the S2M motif and 8-mer region have been deleted to simplify folding calculations (see methods). The predicted secondary structure and thermodynamic stability of this model stem loop and variations are shown. The positions of variations are marked in red.

**Supp. Figure 3.**
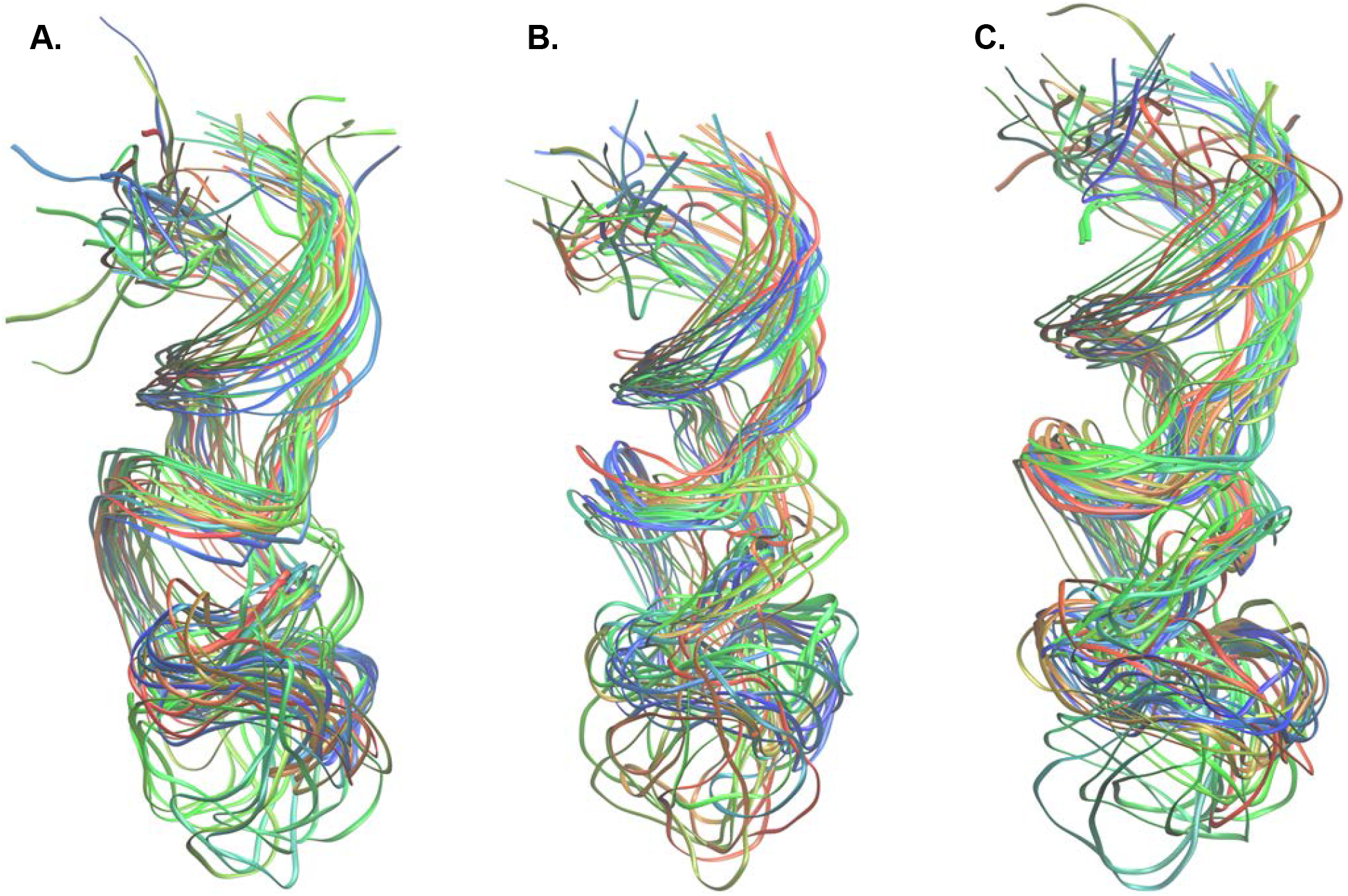
Overlay of the structures from the MD trajectories of the SARS-CoV S2M loop (**A**), G29734C (**B**) and G29742U (**C**) variants. The overlay includes structures from all four 180 ns trajectories collected for each model. Twenty-six structures were selected at equally spaced intervals from the combined trajectories and are colored as a function of time and trajectory from blue to red.

